# Genome scaffolding with PE-contaminated mate-pair libraries

**DOI:** 10.1101/025650

**Authors:** Kristoffer Sahlin, Rayan Chikhi, Lars Arvestad

## Abstract

Scaffolding is often an essential step in a genome assembly process, in which contigs are ordered and oriented using read pairs from a combination of paired-ends libraries and longer-range mate-pair libraries. Although a simple idea, scaffolding is unfortunately hard to get right in practice. One source of problem is so-called PE-contamination in mate-pair libraries, in which a non-negligible fraction of the read pairs get the wrong orientation and a much smaller insert size than what is expected. This contamination has been discussed in previous work on integrated scaffolders in end-to-end assemblers such as Allpaths-LG and MaSuRCA but the methods relies on the fact that the orientation is observable, *e.g.,* by finding the junction adapter sequence in the reads. This is not always the case, making orientation and insert size of a read pair stochastic. Furthermore, work on modeling PE-contamination has so far been disregarded in stand-alone scaffolders and the effect that PE-contamination has on scaffolding quality has not been examined before.

We have addressed PE-contamination in an update of our scaffolder BESST. We formulate the problem as an Integer Linear Program (ILP) and use characteristics of the problem, such as contig lengths and insert size, to efficiently solve the ILP using a linear amount (with respect to the number of contigs) of Linear Programs. Our results show significant improvement over both integrated and standalone scaffolders. The impact of modeling PE-contamination is quantified by comparison with the previous BESST model. We also show how other scaffolders are vulnerable to PE-contaminated libraries, resulting in increased number of misassemblies, more conservative scaffolding, and inflated assembly sizes.

The model is implemented in BESST. Source code and usage instructions are found at https://github.com/ksahlin/BESST. BESST can also be downloaded using PyPI.

## 1. Introduction

Genome assembly is still a challenging process, especially for large genomes, and scientists experiment with different combinations of data and tools to reduce errors, improve contiguity, and avoid ambiguity. An important step in the assembly process is scaffolding, in which contigs are ordered and oriented, and joined to form a larger scaffold unit. The input is a set of contigs and one or several genome mappings of paired short reads from either paired-end (PE) sequencing or mate-pair (MP) sequencing.

Evaluations (*e.g.,* (Hunt et al., 2014)) have shown that scaffolders make many mistakes, perhaps more than one might expect from what could appear a straightforward computational problem. The input to a scaffolder is however both large and noisy, and the data characteristics can vary a lot depending on the organism and assembler. Although contiguity and errors are the most important metrics to evaluate a scaffolder by, we came to note that there are other artifacts from scaffolders not reported in these metrics. We observed that assemblies could increase in size with up to 106% after scaffolding and this mostly affects fragmented assemblies. We call this effect *assembly inflation.* A successful scaffolding will have some assembly inflation due to, *e.g.,* unsequenced regions, but it should in general be very small.

The type of technology, PE or MP, and their main parameter, the insert size, determine how far apart the reads are distributed on the genome and thereby at what distances contigs can be connected into scaffolds. Whether it is PE or MP also determines if the reads are read towards (PE) or apart (MP) from each other. There are numerous complications with PE and MP reads. For example, the insert size is not perfectly controlled and larger insert size typically means a larger variance in the distribution of distances between the reads, making scaffolding harder (Sahlin et al., 2014). A largely ignored problem (on the computational side) is so-called PE contamination of MP libraries, which is a consequence of the MP library preparation. During the process, an unknown fraction of fragments that do not contain the circularization junction are sequenced. These misreads behave like PE reads, with opposite read direction from MP and effectively with a much smaller insert size (Illumina, 2012). Hence, PE contamination reads may confuse a scaffolder that assumes an MP library is clean from contamination, suggesting a different relative order of contigs.

When designing a scaffolder, one can take the stance that PE contamination is (1) an experimental issue which should be controlled in the wet-lab and (2) it can be treated as noise in the MP that will get filtered away in an scaffolding optimization procedure anyway. Regarding (1), we probably have to accept PE contamination as a largely unavoidable difficulty and, in that case, we argue that the PE should be explicitly modeled in MP datasets to reduce errors. Although a decent MP library will contain more true MP reads than PE, and hence overshadow PE contaminants in many cases, assumption (2) will not hold for fragmented assemblies as there are many short contigs close to each other making PE links dominate MP links. The ambition can also be set higher: perhaps we can start utilizing PE contamination as valuable short-range information rather than nuisance that needs to be filtered away?

One may also ask how sensitive the current generation of stand-alone scaffolders are to MP libraries with a high amount of PE contamination? If scaffolders have been designed for near ideal datasets, what can one expect in a more difficult situation?

PE-contamination has been discussed in work on integrated scaffolders in end-to-end assemblers such as **ALLPATHS-LG** (Gnerre et al., 2011) and MaSuRCA (Zimin et al., 2013). These methods relies on the fact that the orientation is observable, *e.g.*, by finding and removing the adapter sequence. These methods are not ideal as a fraction of read pairs will not contain the adapter (depending on shearing size), thus orientation will not be observable. An additional benefit of not assuming orientation is observable is that it does not invoke a constraint on having a sufficiently small fragment shearing size such that one of the reads contains the adapter. when choosing shearing size of the circularized fragments in the library construction protocol. Smaller shearing size gives a higher quantity of “identifiable” mate-pairs (adapter is present) together with higher amounts of chimeric reads and less informative sequence in the mate-pairs (due to adapter present and overlapping reads). On the other hand, shearing size gives less “identifiable” mate-pairs but higher amounts of true mate pairs, and less PE-fragments as well as chimeric reads — due to the increased probability to sequence both ends of a larger fragments on both sides of the adapter. A benefit for scaffolding with larger fragments is that span coverage of the PE-contamination gets better — giving better short range connection.

As this contamination has neither been modeled nor examined in terms of how it degrades scaffolding results, we decided to investigate this. We developed a scaffolder that models PE contamination reads in the scaffolding process on MP libraries. We show that modeling PE-contamination improves scaffolding. Our design utilizes an integer linear program (ILP) for the MP/PE classification and uses information from the interval structure of the contig graph to heuristically obtain a set of linear programs (LP) to solve. The heuristic method is very fast and finds the correct order of contigs in all cases given the model assumptions, as seen on our simulated data. With the new implementation, BESST-v2 completes scaffolding of an assembly for the 20 Gbp spruce genome with five libraries in 24 hours (mapping time excluded).

The new method is benchmarked in a large evaluation over a range of different assembly instances. With the use of PE contamination reads, the new version of BESST places more contigs in fragmented assemblies due to the extra short range information. To account for PE contamination in the scaffolding step is especially important for larger genomes where assemblies tends to be fragmented. This is seen from the significant reduction of errors in BESST-v2 over BESST on the more fragmented assemblies in our evaluation data (Supplemental Table S13).

## 2 Methods

The presented work builds on BESST scaffolder (Sahlin et al., 2014), which iterates over PE and/or MP libraries in the order of their mean insert size. For each library, BESST scaffolds in two steps. First, large contigs are scaffolded using a greedy statistical scoring heuristic (a contig is large if it is unlikely that a read-pair can span over the contig). In a second step, smaller contigs are placed within gaps of the larger contigs, or between larger contigs that were not linked in the first step (*e.g.*, because their distance was larger than the insert size). The second step is done with a breadth-first search where the highest scoring paths are selected. Each path is a local subset of the contig graph, thus naturally dividing the contig graph into subregions in which the contigs needs to be oriented, ordered, and positioned accurately (deciding distance between contigs in a scaffold). If PE contamination is present, it can confuse the ordering as PE reads have the opposite orientation to MP reads. We will here define the problem of ordering and positioning of contigs as an ILP and use heuristic permutation of the contig ordering to efficiently find an assignment with good objective value. Contig orientation will be induced by a chosen path, see section 2.2. Figure 1 shows the possible orderings for two contigs joined by a paired read link. For *m* contigs linked with paired reads, we have *m*! possible orderings and it is not feasible to try them all. However, if we can chose a subset of these orderings to try, we can evaluate each given ordering by finding optimal gap sizes between the contigs. A given ordering is evaluated with a straightforward LP formulation using the gap model by Sahlin et al. (2012).

**Figure 1:**
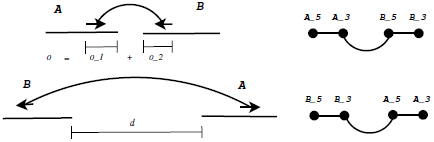
Left: Two possible placements of contigs if we do not know the relative orientation the read pair link. Right: The corresponding contig graphs where “5” and “3” denotes the 5′ and 3′ ends.

### 2.1 Integer Linear Program formulation

The following paragraphs introduce the ILP formulation. First we consider a fixed order of contigs linked by only MP links. We then add PE links to the formulation via an unknown variable representing the order of contigs.

#### Fixed ordering objective without contamination

Let a contig graph G be an undirected graph created by contigs and read pair links where each contig is represented by two vertices (for the 5′and 3′end respectively) and an intra-contig edge. Let an inter-contig edge be an edge in G that connects two vertices that comes from different contigs. An inter-contig edge is created between two vertices if one or more read pairs suggests two contigs and the read pairs (if more than one) suggests the same relative orientation and distance of contigs (known as “link-bundling", Huson et al. (2002)). The intra-contig edges are used in the implementation but does not contribute to the ILP problem and we will from now on only discuss inter-contig edges. Notice that a read pair can give rise to two different edges in G, depending on if the read pair is assumed to be in PE or MP orientation, see Figure 1.

Assume that we have a connected subgraph of G induced by contigs *c*_1_,…, *c*_*m*_ and a set E of inter-contig edges (not all contigs need to have edges between them), see Figure 2 for an example of a contig region. A linear order of *m* contigs has *m* — 1 gaps *g*_*l*_,... *g*_*m-1*_. Let us focus on the edge e between contigs *c_i_* and *c*_*j*_, defined by ***ω**_e_* links. Using only the links between *c_i_* and *c*_*j*_, assume that we have an estimate 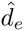 of the distance between these two contigs, see Figure 1. Note that this distance is not to be confused with the gaps *g_i_*,… *g_m-i_*, as *c_i_* and *c_j_* might not be adjacent in the given ordering. How to get the estimate of 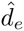 accurately is described in (Sahlin et al., 2012). From 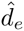 (see Figure 1), we naturally get the average insert size of the links that spans *c_i_* and *c_j_* as 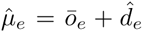, where 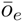 is the average observation. That is 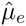 is the average insert size suggested by the links between *c_i_* and *c_j_*. Notice that we define the insert size to include the read lengths (sometimes denoted the fragment length). In a given placement of the m contigs, *c_i_* and *c_j_* will be at distance 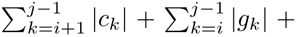, adding up the *j — i —* 1 contigs and *j — i* gaps between *c*_*i*_ and *c*_*j*_. The objective is to minimize the discrepancy between the placement in the current ordering and 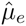, that is, to minimize

**Figure 2:**
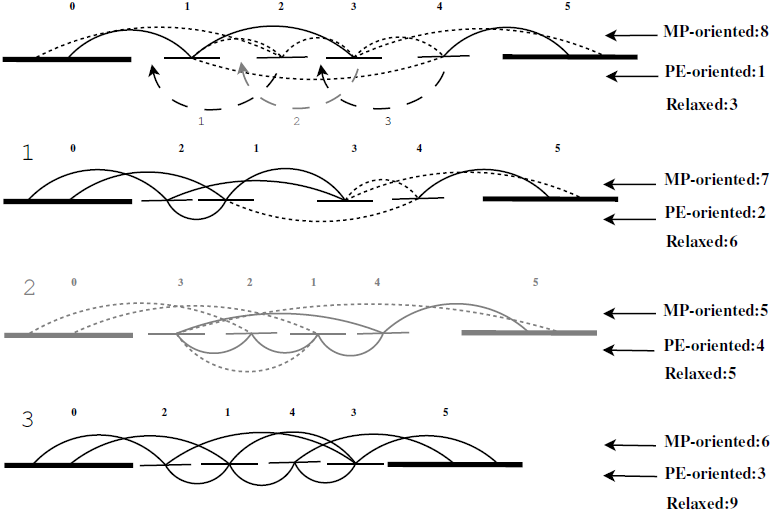
Example of six contigs in an ILP problem of which two of them are border contigs. Edges with MP orientation in a given ordering appears above the contigs, and edges with PE orientation appears below the contigs. An edge appears dotted if it is stretched or compressed in the current solution relative to the predicted distance, *i.e.* 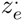 is large. If it has approximately the predicted distance, it appears whole. Notice that the binary states of dotted and whole edges in this figure is only for illustration purposes. In each LP, the value of 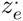 is continuous. Solving the LP for the initial ordering *p*_*0*_ of contigs results in many stretched or compressed edges. After step 1 (moving contig 2) ξ_0_ < ξ_1_, hence p_1_ is accepted (illustrated in figure as links appearing less stretched). Step 2 is rejected as ξ_1_ < ξ_2_ (illustrated as more stretched links than in *p*_*1*_. Finally, the ordering of *p*_*2*_ is accepted as *p*_*2*_ < *p*_*1*_. Note that, The permutation in step 3 will always be to put contig 4 right before contig 3 (index with respect to p_0_), but it is performed on the current lowest objective path, *i.e. p*_1_.

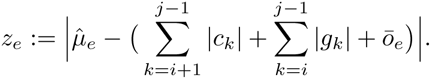

Since each edge have ***ω**_e_* links, we would like to minimize

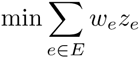

taken over the possible orderings.

#### Fixed ordering objective with contamination

Let 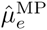 denote the estimated average insert size of links from edge e between two contigs given that the links are oriented as mate pairs in a given placement, and similarly let 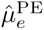 be the estimated average paired end insert size of edge *e*. Analogously, define the distance discrepancies 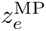 and 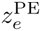 and let **σ**^*PE*^ and **σ**^*MP*^ be the standard deviations of the MP and PE distributions. Let *p* be the proportion of **PE** contamination. We need to adjust for the “observation frequency” between PE- and MP’s in the objective function. Also, larger uncertainty **σ** increases uncertainty in estimations 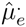, thus we need to weight the penalty in the objective function with this quantity. We have the objective function as

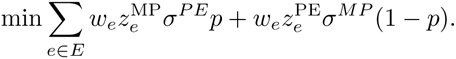

As seen in the objective, edges with larger weights w will penalize a possible distance discrepancy more. Also, as PE links are on average observed with frequency *p*, the distance discrepancy 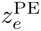 is weighted with (1 — *p*). This is in order to give more weight to the less frequent links, thus balancing the weights that the edges carry. Similarly, wider distributions has larger **σ**. To balance the penalty (allowing larger variation around predictions with high uncertainty) we penalize discrepancies from predicted MP distances with 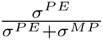 and discrepancies from predicted PE distances with 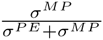, but the denominator is just a scalar and is omitted in the objective function stated above. Note that the model is not well defined if there is no PE contamination and we do not use it if the estimated contamination level is less than 1%.

#### Fixed ordering constraints

Due to the MP library insert size, we do not allow gaps larger than **μ**^MP^ + *k***σ**^*MP*^, with *k* empirically set to 2. Therefore, we have *m* — 1 constraints

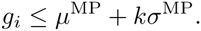

Notice that even though a gap is an integer value, we choose to work with a relaxed problem to be able to apply LP.

#### Formulating an ILP for unknown orientation and ordering

The unknown in this problem is the orientation of contigs in a region. Let 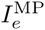 be the indicator function for edge *e* having MP orientation. The full ILP has the variables *g, z* and *I*^MP^ and is written as

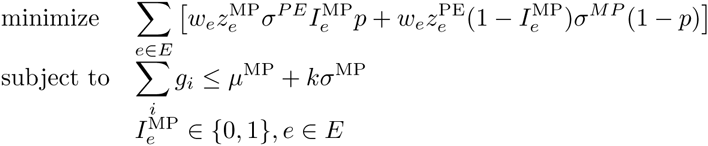

We can use the minimum objective value to this ILP to evaluate contig orderings. For *m* contigs, there are *m*! possible orderings, thus we need a way to efficiently choose a subset of orderings to run the ILP on.

For each assignment of 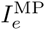, *e*∊E, the problem is a regular LP (remember the assumption g∊R) and we use common LP methods to express the LP on standard form, *i.e.,* introducing help variables to remove absolute values and negative valued variables. This implies that we have 2(*m* —1) + |*E*| constraints in practice. The |*E*| extra constraints comes from the fact that *z*_*e*_ is an absolute value and 2(*m* — 1) constraints because the gap variables are allowed to be negative (letting *g_i_* = *x*_*i*_-*y_i_*, for *x*_*i*_,*y*_*i*_= 0). We solve the LP with the simplex method.

### 2.2 Solving the ILP

We make use of the library information to heuristically and efficiently traverse the space of possible solutions that the ILP permits. The procedure is illustrated in Figure 2. Every region has two “large” border contigs (defined as in (Sahlin et al., 2014)) that are not permuted. The breadth-first search in BESST generates paths in G where a *path* corresponds to a set of contigs where each consecutive pair of contigs is connected by a link with MP orientation (given the initial ordering). The problem is to infer if the links are correctly oriented (MPs) or if they are from PE contamination. Let *p*_*0*_ = *c*_1_,…,*c*_*m*_ be the initial path found by the BFS (excluding the border contigs *c*_*0*_ and *c*_*m*_ + 1) and let the objective value of *p*_*0*_ be ξ_0_. We then iteratively permute *p*_*0*_, in iteration *i***∊**[1, *m* — 1] forming *p*_*i*_ from *p*_*i*-1_ by placing contig *i* + 1 before *i* (making the link from *i* to *i* + 1 PE oriented). If the objective value ξ_i_ < ξ_i-1_ we keep the permutation, otherwise *p*_*i*_ := *p*_*i-1*_ (the permutation is denied). By definition, *p*_*m*-1_ will be the path with lowest objective (of the tried paths). Notice that only consecutive contigs (with respect to *po*) are permuted in this procedure *i.e., i* with *i* + 1, which correspond to testing whether neighboring contigs are linked by MP or PE links. However, the permutation of *i* with *i* + 1 are done with respect to the current lowest objective path *p*_*i*_, in which they might not be neighbors anymore due to previous permutations, see Figure 2.

This heuristic procedure allows us to solve *m* —1 LP’s in a region with *m* contigs. Notice that changing relative orientation of contigs would yield pairs of orientation FF or RR, which are invalid according to our model, thus no such operation are allowed. A solution to each LP gives a set of real valued gaps *g_i_*, *i* ξ [1, *m* — 1] that is used to place the contigs accurately as information from several edges are used simultaneously for each gap.

The model is implemented in the workflow of BESST, and after applying BESST’s scaffolding procedure (Sahlin et al., 2014), we have M ranked clusters of contigs (created by splitting the contig graph based on longer contigs) where the rank is based on how many supporting versus contradicting links a cluster has. Here, we start solving M ILPs, starting from the highest ranked one first. As described, each ILP *i* is solved by solving *m*_*i*_ — 1 LPs, where *m*_*i*_ is the size of ILP *i* **∊** [1, M].

#### Short PE insert size

The success of the limited number of permutations is based on the assumption that the PE distribution is either relatively short, or narrow around the mean. If the PE distribution both is wide and large, it could link both adjacent and more distant contigs, making the graph structure formed by only the true PE reads dense. Many permutations would be needed to untangle the true order of the contigs in such a structure (all *m*! orderings are possible in a clique-like structure). Thus, the permutation of adjacent contigs is efficient if most PE read pairs link adjacent contigs in the true ordering. Although PE reads can span over contigs in an assembly, the model is only affected if many PE links span over contigs (giving large weights to *z*_*e*_). It is our assumption that this is rare compared to adjacent contigs linked by PE reads.

## 3 Results and discussion

We have compared our new implementation of BESST, called BESST-v2 below, with SSPACE (Boetzer et al., 2011), OPERA (Gao et al., 2011), SOPRA (Dayarian et al., 2010), SCARPA (Donmez and Brudno, 2013) and SCAFFMATCH (Mandric and Zelikovsky, 2015). Version 1 of BESST (Sahlin et al., 2014) is included for reference. We also include integrated scaffolder results provided by GAGE, labeled “INTEGRATED".

SSPACE, SCARPA, SOPRA and OPERA were run with mappings from Bowtie (Langmead et al., 2009) as suggested by Hunt et al. (2014) when comparing the aligners BWA (Li and Durbin, 2010), Bowtie and Bowtie2 Langmead and Salzberg (2012). SCAFFMATCH is coupled with Bowtie2. BESST was run with BWA-MEMLi (2013). Full running instructions and details on resource usage are given in Supplementary data.

### 3.1 Datasets

We have included a simulated data set, *sim,* which we developed the model on. First, a set of 354 random contigs was generated, each contig of size 5000 bp (probability 0.2) or 500 bp (probability 0.8). This gave 71 larger contigs and 283 smaller contigs. The largest stretch of smaller contigs consisted of 20 consecutive small contigs. The reference genome was created by concatenating these contigs, hence being of size 496500bp. Based on the reference genome, a MP library with 50x coverage of 2x100bp reads was simulated. The MP library had insert-size distribution N(3000,300) and 30% of the reads were PE contamination with insert size chosen from N(400,40). The larger contigs can be seen as “border contigs” that breaks the scaffolds into separate regions of smaller contigs, because there are no read pairs that can stretch over a large contig. The random selection of contigs ensure different complexities on the subproblems.

Simulated data from the Assemblathon 1 study (Earl et al., 2011) is denoted *assemblathonSk.* We assembled contigs with Minia (Chikhi and Rizk, 2013) from the paired-end reads provided by GAGE (Salzberg et al., 2012). This gave approximately 74000 contigs. The contigs were scaffolded with the 3kbp MP library provided by Assemblathon 1. It contains mate pairs with insert-size distribution N(3000, 300) and 20% PE contamination with an insert-size distribution of N(500, 50).

Three cases, *Staphylococcus aureus, Rhodobacter sphaeroides* and human chromosome 14, were taken from the Genome Assembly Gold-standard Evaluation (GAGE) study (Salzberg et al., 2012) — a comprehensive evaluation of large-scale genome assembly algorithms. We denote them *staph*, *rhodo*, and *hs14* and GAGE provides 8, 9, and 9 contig assemblies, respectively, for these cases. These assemblies were scaffolded with the original shortjump libraries provided by GAGE. By aligning these libraries to the reference genomes with BWA-MEM (Li, 2013) we detected natural PE contamination in the rhodo and hs14 libraries of 41% and 33% respectively with mean insert size of 211 bp and 200 bp respectively (see Supplemental Figures S2 and S3). The staph library had almost no PE contamination with the given alignments (0.0004%, see Figure S1).

We created additional libraries for each reference genome to study the effect of increased levels of PE contamination and PE span coverage. The parameter *c*_*added*_ is introduced to indicate the percentage of added PE reads in a library. The original GAGE libraries have *c*_*added*_ = 0 (0% added contamination). The *c*_*added*_ = 15 libraries were formed by simulating PE reads from the genomes with distribution N(300, 30) and adding them to the original read library. Similarly, the *c*_*added*_ = 40 libraries were formed by simulating PE reads from the genomes with distribution N(400, 40) and adding them to the original library.

Two additional real MP libraries, one for *Rhodobacter sphaeroides* and one for *Escherichia coli,* taken from the study by Ribeiro et al. (2012), were also used in the evaluation. The analysis is found in Supplementary data Section 6.

### 3.2 Evaluation method

Assembly evaluation is known to be difficult (Earl et al., 2011; Salzberg et al., 2012; Hunt et al., 2014), as there are many metrics to consider. Ultimately, the end result of an assembly should be as long error-free sequences as possible. However, two assemblies with the same lengths on error free sequences can still differ if one of the assemblers/scaffolders contains false joins, making the assembly look more contiguous. Thus, errors is another important metric. We choose to look at the adjusted E-size (the expected length of error free sequence in the assembly, Salzberg et al., 2012) which we denote by *E′,* and the number of errors.

Moreover, we need an informative way to present the quality increase/decrease of an assembly from contigs to scaffolds. Since the GAGE data sets contains assemblies of varying quality (with respect to contig errors and *E′*) of each organism, we present the quality improvement between contigs and scaffolds. Let *E′_c_* and *E′_s_* denote the adjusted E-size of the original contig assembly and the scaffolded assembly. We report the increase in errors from original contigs to scaffolds as well as the ratio *E′_s_/E*’_c_. Tables S2-S18 in Supplementary data contains results for individual experiments and the summary tables presented here shows the average increase on each organism and library.

#### Assembly size inflation

As another aspect of scaffolding quality, we also show how total assembly size grows after scaffolding. We observed that some scaffolders increase the assembly size more than others and we wanted to find the cause. As no scaffolder (in this comparison) adds contig sequence on multiple places (*e.g.*, repeats) or derive new sequence from the reads, inflation is due to added gaps (stretches of N’s). We categorize inflation into three possible causes: (1) The genomic content between two contigs is not present in the assembly, thus an approximately correct number N’s are inserted. This is a correct behaviour from a scaffolder and, to some extent, it improves the quality of the assembled genome. (2) The genomic content between two contigs *a* and *c* is present in another contig *b* in the assembly, but the scaffolder is unable to place b. The scaffolder creates (*a* correct number of) N’s between *a* and *c* and leaves *b* isolated in the scaffold Fasta file. This increases the assembly size, as there are two versions of a single location in the assembly, represented by b and N’s respectively. It is not a misassembly, but it can confuse downstream analysis, and it negatively affects the assembly quality, such as spurious size inflation. (3) Errors: a gap between contigs is inserted due to a false join, or the gap sequence significantly differs in length to the real gap length. This is seen as a misassembly that decreases the quality of the assembly. When we discuss assembly inflation we will refer to these three causes as (1), (2) or (3). We also report the ratio of scaffolded assembly size compared to initial contig size, labeled *inflation* in the tables, and find that some scaffolders are prone to (2).

#### Note on evaluation

Snakemake (Koster and Rahmann, 2012) was used to run our evaluation pipeline. QUAST (Gurevich et al., 2013) was used for evaluating the scaffolders as it uses alignments to a reference to identify misassembly breakpoints. Support for computing *E*-size was added. QUAST classifies a misassembly as a breakpoint in the scaffold where the left and right flanking bases differs more than N base pairs from the reference sequence. We have set N = 100 to allow for reasonable variation in gap size estimations.

QUAST does not handle allele shifting of contigs on scaffolds from a diploid genome with a diploid reference. As the assemblathon3k dataset provides both copies of each chromosome, the evaluation was performed by giving only one of the copies of each chromosome (copy A) as reference to QUAST. Therefore, some of the errors occur due to the omitted reference copy. For example, the original assembly of Minia is free from misassemblies if both copies is given as references. The scenario is however the same for all scaffolders so relative performance can still be compared.

#### Note on testing

It is of general importance to know what data has been used in the development of the algorithm. By modifying the ILP, we can obtain something that works well for our data but does not generalize to other data sets. We used simulated data sets similar to the dataset “sim” as problem instances for BESST-v2 when developing the ILP. Our method is extremely accurate for all contamination levels and contig sizes that we have tested. Therefore, it is not strange that we excel on the data set “sim” and BESST’s results on this data set should be seen as a test of correctness in the implementation and formulation of the ILP, given its assumptions. However, these datasets also illustrate how vulnerable other scaffolders can be to paired end contamination in such scenarios. The Assemblathon 1 dataset with Minia’s contig assembly was the data set where we discovered that paired end contamination affected BESST’s result, and we ran BESST before and after the implementation to verify the improvement on this data set that more closely mimics a biological data set. The rest of the data sets were not scaffolded before the model described above was developed.

### 3.3 Inflated assembly sizes

In general, all scaffolders increase the assembly size to some extent but there is large variation among them. On the sim data set, see Table 1, all scaffolders except BESST-v2 and SCARPA inflates the assembly size between 18-106.6% even though all contigs has the potential to be linked (as done in BESST-v2). SCAFFMATCH inflates the assembly size the most (106.6%) and manual inspection reveals that this is due to a mix of cause (2) and (3). SCAFFMATCH uses an approach (maximum matching) that only permits one neighbour of every contig to be joined; this is a limitation on assemblies where the insert size is relatively large compared to contigs, making mate pairs link to several neighbors. SCAFFMATCH addresses this limitation by including a separate insertion-step that attempts to place remaining singleton contigs, but it does not perform well on this data set. SSPACE also inflates the assembly size of the dataset sim to a large extent, but has only one error. The inflation is from (2), which is a result of SSPACE′s heuristic; SSPACE extends scaffolds greedily by choosing the neighboring contig with the most links and may therefore miss many small intermediate contigs. OPERA, SOPRA and BESST shows vulnerability by making orientation mistakes (as in Figure 1) due to creating many joins where the link is interpreted as a MP instead of a PE. SCARPA has low inflation on this data set.

**Table 1:** Results on the sim dataset. BESST-v2 finds the correct order, orientation and approximate positions of all contigs and joins the contigs into a single scaffold. SSPACE gives only one error but places very few contigs and has an inflation of 43.5% from type (2). OPERA and BESST are the most sensitive to PE contamination with respect to misassemblies. “contigs” denotes the initial contig assembly metrics on which the relative in inflation, increase in errors and corrected contiguity are computed.

| Tool | Infl. | Scf-errors | $E'_s/E'_c$ |
| --- | --- | --- | --- |
| BESST-v2 | 1.002 | **0** | **130.9** |
| BESST | 1.370 | 199 | 2.4 |
| SSPACE | 1.435 | 1 | 5.5 |
| OPERA | 1.406 | 255 | 1.0 |
| SOPRA | 1.179 | 73 | 2.0 |
| SCARPA | 1.015 | 9 | 5.4 |
| SCAFFMATCH | 2.066 | 95 | 2.5 |
| | Size | Ctg-errors | $E'_c$ |
| contigs | 496500 | 0 | 3692.0 |

On the assemblathon3k dataset, see Table 2, BESST-v2 is able to increase the adjusted E-size with a factor of 22 while keeping the size of the assembly relatively constant. SCAFFMATCH, SSPACE, BESST and SCARPA inflates the assembly size significantly, but the large amount of errors and small *E’_s_/E′_C_* (for SSPACE and SCAFFMATCH) suggests that this is due to cause (3). As the PE oriented read has mean insert size of 500, this is the data set where PE contamination has the highest span coverage and therefore most likely poses the biggest challenge.

**Table 2:** Assemblathon3k, the original Assemblathon 1 data set with 3kbp short jump library. BESST-v2 removes 83.6% of BESST’s errors and has superior contiguity and number of misassemblies to the other scaffolders. SOPRA is missing from the table because it did not meet the runtime constraint. “contigs” denotes the initial contig assembly metrics on which the relative in inflation, increase in errors and corrected contiguity are computed.

| Tool | Infl. | Scf-errors | $E'_s/E'_c$ |
| --- | --- | --- | --- |
| BESST-v2 | 1.015 | **2167** | **21.9** |
| BESST | 1.137 | 13229 | 4.6 |
| SSPACE | 1.110 | 26410 | 1.0 |
| OPERA | 1.015 | 10161 | 1.7 |
| SCARPA | 1.070 | 15038 | 1.2 |
| SCAFFMATCH | 1.200 | 34124 | 1.0 |
| | Size | Ctg-errors | $E'_c$ |
| contigs | 120105427 | 7 | 5374.9 |

On the GAGE datasets (Tables 3-5) the inflation levels vary significantly among scaffolders. This phenomenon is most evident for SCAFFMATCH and SSPACE and is likely an artifact of their methodology. SCAFFMATCH inflates the assembly size the most and shows larger inflation as contamination increases (up to 19.2% and 28.7% on average on rhodo and hs14 respectively). SSPACE also shows high inflation rate on the GAGE assemblies already with the original libraries (*c_added_* = 0), *e.g.*, 6% on hs14. The contamination further worsens this behavior for SSPACE to 17% on average on rhodo and 9% on hs14 for *c_added_* = 40. SCARPA and BESST are also affected by assembly inflation, but not to the same extent as SSPACE and SCAFFMATCH. Notably, SCAFFMATCH, SSPACE and SCARPA show an extreme inflation in assembly size on the two most fragmented assemblies on rhodo (ABySS and SGA) with inflation between 28-49% on ABYSS and 58-103% on SGA (Supplemental data, Table S7-S9), and a similar trend for the most fragmented assemblies on hs14 (Supplemental data, Table S11-S13). Such extreme inflation is clearly not correct as it almost doubles the assembly size, especially when the contig assembly size is already larger than the true genome size. We also argue that the more subtle inflation on the higher quality data sets are artifacts as BESST-v2 in general has the fewest errors and highest increase in contiguity with only a small increase in assembly size. Inflation/deflation level vary among the integrated scaffolders. For example, SGA contig assemblies is generally significantly larger than the genome size and SGA’s scaffolder removes a lot of sequence in the scaffolding step. In the hs14 assemblies, the integrated scaffolders in Velvet, Bambus2, MSR-CA and SOAPdenovo inflates the assembly size the most and also contains significantly more errors than the other scaffolders.

#### Summary inflation

In total, our results indicate that assembly inflation is more likely due to poor scaffolding (cause (3)) or the inability to place many contigs in a fragmented scaffold (cause (2)) rather than correctly added sequence gaps (1) for both stand alone and integrated scaffolders. This is supported by investigating the individual assemblies (Tables S2-S14). The more fragmented assemblies show significantly higher inflation and error rate than the higher quality ones. We suggest users of scaffolders to look at similar metrics after the scaffolding step is performed.

Notably, BESST-v2 reduces the assembly size slightly with increased PE contamination (Tables 3-5). This is due to the fact that BESST-v2 can use the extra short range information to place smaller contigs, thus lowering inflation by reducing gapped sequence, *i.e.,* gaps caused by (2).

### 3.4 Errors and contiguity

The sim and assemblathon3k datasets show how strongly PE contamination can affect scaffolding (Tables 1-2). There are extreme differences in inflation, errors, and *E′_S_* among the scaffolders on these two datasets. The differences in result between BESST and BESST-v2, as well as the number of errors of, *e.g.,* OPERA, SOPRA and SCAFFMATCH, indicate that a large part of the links created in the scaffolding graph are from PE contamination. BESST-v2 corrects all errors on sim and the majority of errors on assemblathon3k, which indicates that it is an efficient method for finding the correct ordering.

The GAGE datasets (Tables 3-5) further show that PE contamination affects scaffolding significantly, especially when assemblies are more fragmented, as is common for more complex genomes (see Table 5). OPERA and BESST seem to be the most vulnerable to contamination with significant increase in errors and lower *E′_S_* across most assemblies as the contamination level increases. SCARPA is another scaffolder where introduced contamination changes the results drastically across single assemblies. However, one assembly (MSR-CA) is missing from SCARPA on hs14 due to our computational time constraint and the average is therefore somewhat distorted for hs14 with *c_added_* = 40. The effect on individual assemblies is an increasing number of misassemblies (see Tables S12-S14).

**Table 3:**
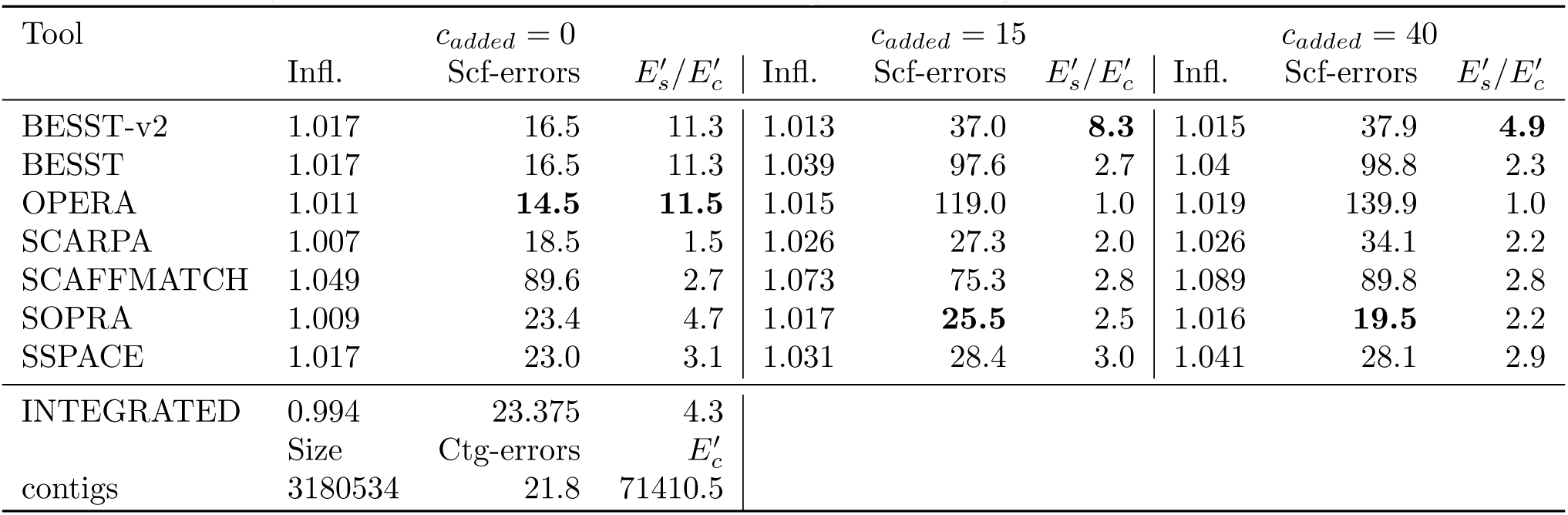
Staph GAGE contig assemblies. Scaffolded with GAGE′s shortjump MP library, *c_added_* = 0, with an added 15% PE contamination reads, N(300, 30), *c_added_* = 15, and with an added 40% PE contamination reads, N(400, 40), *c_added_* = 40. The numbers are averaged over each assembly. Full tables are found in Supplementary data. “contigs” denotes the initial contig assembly metrics on which the relative in inflation, increase in errors and corrected contiguity are computed.

| Tool | $c_{added} = 0$ | | | $c_{added} = 15$ | | | $c_{added} = 40$ | | |
| --- | --- | --- | --- | --- | --- | --- | --- | --- | --- |
| | Infl. | Scf-errors | $E'_s/E'_c$ | Infl. | Scf-errors | $E'_s/E'_c$ | Infl. | Scf-errors | $E'_s/E'_c$ |
| BESST-v2 | 1.017 | 16.5 | 11.3 | 1.013 | 37.0 | **8.3** | 1.015 | 37.9 | **4.9** |
| BESST | 1.017 | 16.5 | 11.3 | 1.039 | 97.6 | 2.7 | 1.04 | 98.8 | 2.3 |
| OPERA | 1.011 | **14.5** | **11.5** | 1.015 | 119.0 | 1.0 | 1.019 | 139.9 | 1.0 |
| SCARPA | 1.007 | 18.5 | 1.5 | 1.026 | 27.3 | 2.0 | 1.026 | 34.1 | 2.2 |
| SCAFFMATCH | 1.049 | 89.6 | 2.7 | 1.073 | 75.3 | 2.8 | 1.089 | 89.8 | 2.8 |
| SOPRA | 1.009 | 23.4 | 4.7 | 1.017 | **25.5** | 2.5 | 1.016 | **19.5** | 2.2 |
| SSPACE | 1.017 | 23.0 | 3.1 | 1.031 | 28.4 | 3.0 | 1.041 | 28.1 | 2.9 |
| INTEGRATED | 0.994 | 23.375 | 4.3 |  |  |  |  |  |  |
| | Size | Ctg-errors | $E'_c$ | | | | | | |
| contigs | 3180534 | 21.8 | 71410.5 |  |  |  |  |  |  |

**Table 4:** Rhodo original GAGE contig assemblies. Sca_olded with GAGE′s shortjump MP library, *c_added_* = 0, with an added 15% PE contamination reads, N(300, 30), *c_added_* = 15, and with an added 40% PE contamination reads, N(400, 40), *c_added_* = 40. The numbers are averaged over each assembly. Full tables are found in Supplementary data. “contigs” denotes the initial contig assembly metrics on which the relative in ination, increase in errors and corrected contiguity are computed.

| Tool | $c_{added} = 0$ | | | $c_{added} = 15$ | | | $c_{added} = 40$ | | |
| --- | --- | --- | --- | --- | --- | --- | --- | --- | --- |
| | Infl. | Scf-errors | $E'_s/E'_c$ | Infl. | Scf-errors | $E'_s/E'_c$ | Infl. | Scf-errors | $E'_s/E'_c$ |
| BESST-v2 | 1.031 | 36.7 | **8.2** | 1.024 | 68.8 | **7.9** | 1.018 | **45.9** | **8.6** |
| BESST | 1.04 | 50.1 | 8.0 | 1.063 | 173.4 | 6.3 | 1.062 | 180.4 | 6.0 |
| OPERA | 1.015 | 146.4 | 2.3 | 1.012 | 206.0 | 1.0 | 1.018 | 439.3 | 1.0 |
| SCARPA | 1.027 | 45.2 | 1.5 | 1.08 | **67.6** | 1.0 | 1.116 | 114.6 | 1.1 |
| SCAFFMATCH | 1.11 | 186.8 | 4.8 | 1.135 | 182.0 | 4.8 | 1.192 | 197.3 | 4.6 |
| SOPRA | 1.018 | 30.7 | 1.6 | 1.071 | 88.9 | 1.5 | 1.115 | 81.1 | 1.4 |
| SSPACE | 1.017 | **25.2** | 1.5 | 1.06 | 76.0 | 1.4 | 1.167 | 155.3 | 1.4 |
| INTEGRATED | 1.033 | 46.444 | 5.367 |  |  |  |  |  |  |
| | Size | Ctg-errors | $E'_c$ | | | | | | |
| contigs | 4640197 | 63.0 | 24662.2 |  |  |  |  |  |  |

**Table 5:** Hs14 original GAGE contig assemblies. Scaffolded with shortjump mate pair reads original library, *c_added_* = 0, with an added 15% PE contamination reads from N(300, 30), *c_added_* = 15, and with an added 40% PE contamination reads from N(400, 40), *c_added_* = 40. The numbers are averaged over each assembly. Full tables are found in Supplementary data. “contigs” denotes the initial contig assembly metrics on which the relative in inflation, increase in errors and corrected contiguity are computed.

| Tool | $c_{added} = 0$ | | | $c_{added} = 15$ | | | $c_{added} = 40$ | | |
| --- | --- | --- | --- | --- | --- | --- | --- | --- | --- |
| | Infl. | Scf-errors | $E'_s/E'_c$ | Infl. | Scf-errors | $E'_s/E'_c$ | Infl. | Scf-errors | $E'_s/E'_c$ |
| BESST-v2 | 1.03 | **2667.8** | **3.7** | 1.028 | **2916.2** | **3.7** | 1.019 | 2337.7 | **3.3** |
| BESST | 1.046 | 4292.4 | 3.0 | 1.057 | 6343.8 | 2.7 | 1.049 | 6015.8 | 2.4 |
| OPERA | 1.024 | 4854.3 | 1.5 | 1.007 | 5587.4 | 1.1 | 1.021 | 7526.9 | 1.0 |
| SCARPA | 1.036 | 2990.7 | 1.6 | 1.046 | 3499.6 | 1.9 | 1.046 | 3490.3 | 1.8 |
| SCAFFMATCH | 1.128 | 9345.3 | 2.3 | 1.132 | 9321.7 | 2.3 | 1.287 | 9687.7 | 2.4 |
| SOPRA | 1.044 | 5116.1 | 2.4 | 1.045 | 4382.7 | 2.2 | 1.011 | **1109.8** | 1.7 |
| SSPACE | 1.063 | 3940.3 | 2.6 | 1.065 | 3913.9 | 2.6 | 1.093 | 3650.8 | 2.5 |
| INTEGRATED | 1.079 | 7238.222 | 2.511 |  |  |  |  |  |  |
| | Size | Ctg-errors | $E'_c$ | | | | | | |
| contigs | 98516181 | 2109.1 | 14087.4 |  |  |  |  |  |  |

SSPACE takes a more conservative approach as it only chooses one neighboring contig to extend the scaffold with, which is reflected with increased level of inflation. This results in a slightly lower *E′_s_* for the *c_added_* = 40 libraries. On rhodo (Table 4), however, there is a large increase in errors, where almost the entire increase is on the two fragmented assemblies ABySS and SGA. Due to the number of failed runs by SOPRA on hs14 (from the time constraint), no general conclusion about trend can be made with the averaged data presented here. However, as with the other scaffolders, SOPRA is the most affected by fragmented assemblies and greatly increases the number of errors on SGA and Velvet on hs14 when contamination is present (Table S12-S14). SCAFFMATCH shows fairly consistent number of misassemblies and *E′_s_* across all datasets and with a relatively good *E′_S_*. However, it always has significantly more misassemblies than other scaffolders.

With the GAGE libraries, BESST-v2 has the second fewest to fewest misassemblies and second highest to highest *E′_S_* for almost all runs. The *E′_S_* even increases on rhodo (Table 4) as contamination is introduced due to the extra read pair information. Although average number of misassemblies increases slightly for BESST-v2 with the contamination level in the GAGE runs, the difference is relatively small. In fact, the number of misassemblies at *c_added_* = 40 made by BESST-v2 are competitive also at *c_added_* = 0 with other scaffolders. The tradeoff between misassemblies and increase in 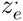 gives BESST-v2 favorable results to the other scaffolders. Table 5 for *c_added_* = 0 shows that BESST-v2 lowers the misassemblies with almost 40%, with the most significant decrease on ABySS, ABySS2, SGA and Velvet. This suggests that a large part of the edges formed in a fragmented contig graph comes from PE contamination. Notably, BESST-v2 also improves scaffolding compared with integrated scaffolder across almost all data sets, Table S4,S7,S12. On hs14, BESST-v2 has both lower amount of errors and higher corrected contiguity for all assemblies except for ABySS and ABySS2, where more errors are introduced but contiguity is 3-fold higher.

In conclusion, stand-alone scaffolders introduce a large number of misassemblies on the fragmented assemblies with contamination present (*e.g.*, ABySS and SGA in rhodo, Table S7-S9, or ABySS, SGA and Velvet in hs14, Table S12-S14). By comparing the two versions of BESST, we see that BESST-v2 corrects most of these misassemblies (see *e.g.,* ABySS and SGA assemblies on rhodo, Table S7-S9, and hs14, Table S12-S14, in Supplementary data). Modeling PE contamination in the scaffolding step is important for larger and more complex genomes where assemblies tends to be fragmented, thus the proportion of PE links increases. This is supported by looking at the number of errors that is corrected in the hs14 assemblies on real data; with original GAGE libraries BESST-v2 reduce 54% of BESST’s errors when scaffolding the SGA assembly and 53% on the ABySS assembly (See Supplemental Table S13). Finally, BESST-v2 generally gives preferable results over integrated scaffolders on GAGE data.

### Runtime and alignments

We also measured runtime and peak memory of the tools. Resource usage is not the main objective in this study and has been studied in other work (Hunt et al., 2014; Sahlin *et al.,* 2014). However, we note that BESST, BESST-v2, SSPACE, OPERA, and SCAFFMATCH (with the greedy approach) all have runtimes that should be practical on most genomes. There is no big difference in speed and memory demand between BESST-v2 and BESST.

## 4 Conclusions

We have designed a scaffolder that can identify and make use of read pairs with PE orientation in a MP library, so called PE contamination. The scaffolder (BESST-v2) accurately infers scaffolds, even with high levels of contamination, and we showed that other scaffolders are vulnerable to PE-contaminated libraries. Our results indicate that, when modeled, PE contamination helps scaffolding, serving as short-range linking information which complements long-ranging mate-pair reads. This combination of reads helps placing small contigs in fragmented assemblies. We also showed that inflated assembly sizes after scaffolding are more often a result of the inability of scaffolders to place all contigs in a scaffold or erroneous gaps, rather than correctly inserted unknown sequence.

